# MdMYB3 helps regulate anthocyanin accumulation in apple calli under moderately acidic conditions

**DOI:** 10.1101/429456

**Authors:** Yi-Cheng Wang, Jing-Jing Sun, Yan-Fen Qiu, Xiao-Jun Gong, Li Ma, Zong-Ying Zhang, Shou-Qian Feng, Yan-Ling Wang, Xue-Sen Chen

**Affiliations:** College of Forestry, Shandong Agricultural University, Tai-An, 271018, Shandong, China; State Key Laboratory of Crop Biology, College of Horticulture Science and Engineering, Shandong Agricultural University, Tai-An, 271018, Shandong, China

**Author notes:** Corresponding authors: Yan-Ling Wang and Xue-Sen Chen (Y.-L. Wang); (X.-S. Chen).

## Abstract

Anthocyanins are the key factors controlling the coloration of plant tissues. However, the molecular mechanism underlying the effects of environmental pH on the synthesis of apple anthocyanins is unclear. In this study, we analyzed the anthocyanin contents of apple calli cultured in media at different pHs (5.5, 6.0, and 6.5). The highest anthocyanin content was observed at pH 6.0. Additionally, the moderately acidic conditions up-regulated the expression of *MdMYB3* as well as specific anthocyanin biosynthesis structural genes (*MdDFR* and *MdUFGT*). Moreover, the anthocyanin content was higher in calli overexpressing *MdMYB3* than in the wild-type controls at different pHs. Yeast one-hybrid assay results indicated that MdMYB3 binds to the *MdDFR* and *MdUFGT* promoters *in vivo*. An analysis of the *MdDFR* and *MdUFGT* promoters revealed multiple MYB-binding sites. Meanwhile, electrophoretic mobility shift assays confirmed that MdMYB3 binds to the *MdDFR* and *MdUFGT* promoters *in vitro*. Furthermore, GUS promoter activity assays suggested that the *MdDFR* and *MdUFGT* promoter activities are enhanced by acidic conditions, and the binding of MdMYB3 may further enhance activity. These results implied that an acid-induced apple MYB transcription factor (MdMYB3) promotes anthocyanin accumulation by up-regulating the expression of *MdDFR* and *MdUFGT* under moderately acidic conditions.

## Introduction

Anthocyanins are water-soluble natural pigments that are widely distributed in plants. They are responsible for the diverse colors observed in plant tissues and organs, and are important antioxidants [1,2]. Their potential antioxidant activities help to protect plants from oxidative damage and have useful health-related benefits for humans [3].

Anthocyanins are synthesized in a distinct branch of the flavonoid biosynthetic pathway that has been well characterized in plants [4]. In addition to the anthocyanin biosynthesis genes, R2R3 MYB transcription factors (TFs) can function cooperatively with basic helix-loop-helix (bHLH) and WD40 TFs to regulate anthocyanin biosynthesis [5–8]. In apple, three R2R3 MYB TFs (MdMYB1, MdMYB10, and MdMYBA) have been functionally characterized and were confirmed to be responsible for anthocyanin accumulation based on their ability to directly regulate the expression of anthocyanin biosynthesis structural genes [9]. Anthocyanin biosynthesis is also stimulated by abiotic factors, including low temperatures, hormones, high irradiance, sugar, and pH [10–14]. There is clear evidence that high pH levels enhance the degradation of anthocyanins in carrot [12], while exposure to low pH induces the expression of anthocyanin biosynthesis genes in crabapple leaves, ultimately resulting in increased anthocyanin levels [15].

Previous studies revealed that the accumulation of anthocyanins in the vacuoles of plant tissue cells depends on the pH of the vacuoles in which anthocyanins localize [16, 17]. In morning glory (*Ipomoea tricolor*) petals, the pH of vacuoles is relatively low when the flower bud opens, causing the petals to be red, but the vacuolar pH increases over time and the petals eventually turn blue [18]. This color change is due to the Na^+^/H^+^ exchanger encoded by the *PURPLE* gene, which transports sodium ions into the vacuole and moves protons out of the vacuole to increase the pH [19]. *Petunia hybrida* flowers normally have a lower pH than *I. tricolor* flowers, and the wild-type (WT) flowers remain on the reddish (low pH) side of the color spectrum. The PH4 gene encodes a MYB domain and is expressed in the epidermis of petals [20]. A mutation to *PH4* results in the lightening of petal colors as well as increases in the pH of petal extracts and, in some genetic backgrounds, the elimination of anthocyanins in flowers [20]. Additionally, mutations to *ANTHOCYANIN1* (*AN1*), *AN2*, and *AN11* lead to a loss of anthocyanin pigments and an increase in the pH of petal extracts [21]. In apple, some MYB TFs control cell pH, and the vacuolar accumulation of anthocyanins and malate is conserved between apple and *Arabidopsis thaliana* [22, 23]. However, there have been relatively few reports describing the effects of environmental pH on the color of apple tissue.

Anthocyanin contents are low and unstable in the flesh of most cultivated apples [24]. *Malus sieversii* f. *niedzwetzkyana* fruits are the red variant of *M. sieversii* fruits. Moreover, *M. sieversii* f. *niedzwetzkyana* branches, leaves, flowers, and fruit peel and pulp are all red because of their very high anthocyanin concentrations, which may provide considerable health benefits for humans [25]. Therefore, investigating the mechanism underlying anthocyanin biosynthesis in *M. sieversii* f. *niedzwetzkyana* is warranted, and may promote the development of new apple varieties that produce fruits with high anthocyanin contents. Additional research may also provide new information about the genetic diversity among cultivated apple species. In this study, we analyzed the effects of moderate pH changes on the accumulation of anthocyanins in red-fleshed apple calli produced from the leaves of an R6/R6 homozygous line, which is the hybrid offspring of a cross between *M. sieversii* f. *niedzwetzkyana* and *Malus domestica* ‘Fuji’. The *MdMYB3* gene was isolated because it was differentially expressed between calli exhibiting good or poor coloration after an exposure to low or high pH levels. The function of this gene during anthocyanin biosynthesis was also characterized by examining the effects of different pH treatments. Therefore, we herein provide new insights into how MYB TFs influence pH-regulated anthocyanin biosynthesis in red-fleshed apple fruits.

## Materials and methods

### Plant materials and pH treatments

A red-fleshed apple callus was induced as previously described [26]. Murashige and Skoog (MS) medium supplemented with 1 mg/L 6-BA and 0.3 mg/L NAA was used for subculturing. The medium pH was adjusted to 5.5, 6.0, or 6.5. Calli were grown in an incubator at 24 °C under a 16-h light/8-h dark photoperiod (light intensity: 1,000–2,000 lx). Calli were harvested and subcultured every 18 days.

### Measurement of anthocyanin content

Total anthocyanins were extracted from each sample according to the methanol-HCl method developed by Ji [26]. The absorbance of the resulting solutions was analyzed at 530 nm using a UV-2450 spectrophotometer (Kyoto, Japan) to calculate anthocyanin levels.

### Quantitative real-time PCR (qRT-PCR) analysis

Total RNA was extracted from calli using the RNAprep Pure Plant Kit (Tiangen, Beijing, China), after which first-strand cDNA was synthesized using the RevertAid First Strand cDNA Synthesis Kit (Fermentas, St. Leon-Roth, Germany). Gene expression levels were analyzed in a qRT-PCR assay, which was completed with 50 ng/μL cDNA as the template, the SYBR Green PCR Master Mix (TransGen Biotech, Beijing, China), and the iCycler iQ5 system (Bio-Rad, Hercules, CA, USA). Each sample was analyzed with three replicates, and the data are presented in bar graphs as the mean ± standard error. Details regarding the qRT-PCR primers are provided in Supplementary Table S1. The MdActin gene was used as the internal control. The data were analyzed by the 2^−ΔΔCt^ method.

### Transformation of the red-fleshed apple callus

Gene-specific primers were designed to prepare a construct for the overexpression of *MdMYB3* (described in Supplementary Table S1). The open reading frame was incorporated into the pRI101-AN vector (Takara, Otsu, Japan), which was then inserted into *Agrobacterium tumefaciens* LBA4404 cells. Two-week-old calli grown in liquid medium were co-cultured with *A. tumefaciens* LBA4404 cells carrying the 35S:MdMYB3-GFP construct on MS medium containing 2 μmol/L NAA and 4 μmol/L 6-BA at 25 °C for 2 days in darkness. The calli were then transferred to fresh MS medium supplemented with 662 μmol/L carbenicillin and 74 μmol/L kanamycin to screen for transformants carrying the transgene.

### Yeast one hybrid (Y1H) analysis

Yeast one-hybrid assays were completed with yeast strain Y187 (Clontech,) according to the manufacturer’s instructions. The MdMYB3 gene was cloned into the pGADT7 vector, while the promoters of anthocyanin biosynthesis structural genes were inserted into separate pHIS2 vectors. Yeast Y187 cells were co-transformed with different vector combinations, after which the resulting interactions were examined on synthetic defined medium lacking Trp, Leu, and His (SD/−Trp/−Leu/−His), but supplemented with an optimal concentration of 3-AT.

### Electrophoretic mobility shift assay

Electrophoretic mobility shift assays (EMSAs) were performed using biotinylated probes and the LightShift Chemiluminescent EMSA kit (Thermo Scientific, Waltham, MA, USA). The *MdMYB3* gene was recombined into the pET32a vector, which was then inserted into *Escherichia coli* BL21 (DE3) cells for the subsequent production of the MdMYB3-His fusion protein. The fusion protein was purified using the Ni-Sepharose His-Tag Protein Purification Kit (CWBiotech, Beijing, China). All probes specific for the promoter fragments were obtained from Sangon Biotechnology Co., Ltd. (Shanghai, China). The EMSA reaction solutions included the following components: 2 μL 1× binding buffer (2.5% glycerol, 50 mmol/L KCl, 5 mmol/L MgCl_3_, and 10 mmol/L EDTA), 10 μL HIS protein or MdMYB3-His fusion protein, and 1 μL probe. The solutions were incubated at room temperature for 25 min. Unlabeled probes served as competitive probes. The EMSAs were conducted as described by Xu [27].

### β-glucuronidase (GUS) analysis

The MdDFR and MdUFGT promoter sequences were integrated into separate pBI121 vectors, which included a sequence encoding GUS. The constructed proMdDFR:GUS and proMdUFGT:GUS vectors were transferred to WT and OE-MdMYB3 calli. The transgenic calli were then immersed in GUS staining buffer and incubated at 37 °C for 10 h. The GUS enzyme activity was analyzed with 0.3 g transgenic callus extracted using GUS buffer. Protein concentrations were determined according to the Bradford method [28]. The GUS enzyme activity was measured as previously described by Xie [29].

## Results

### Effects of pH on callus anthocyanin content

The anthocyanin content of red-fleshed apple calli changed considerably in response to various pH levels (Fig. 1). The highest anthocyanin content was observed at pH 6.0, and resulted in reddish calli (Fig. 1).

**Fig. 1.**
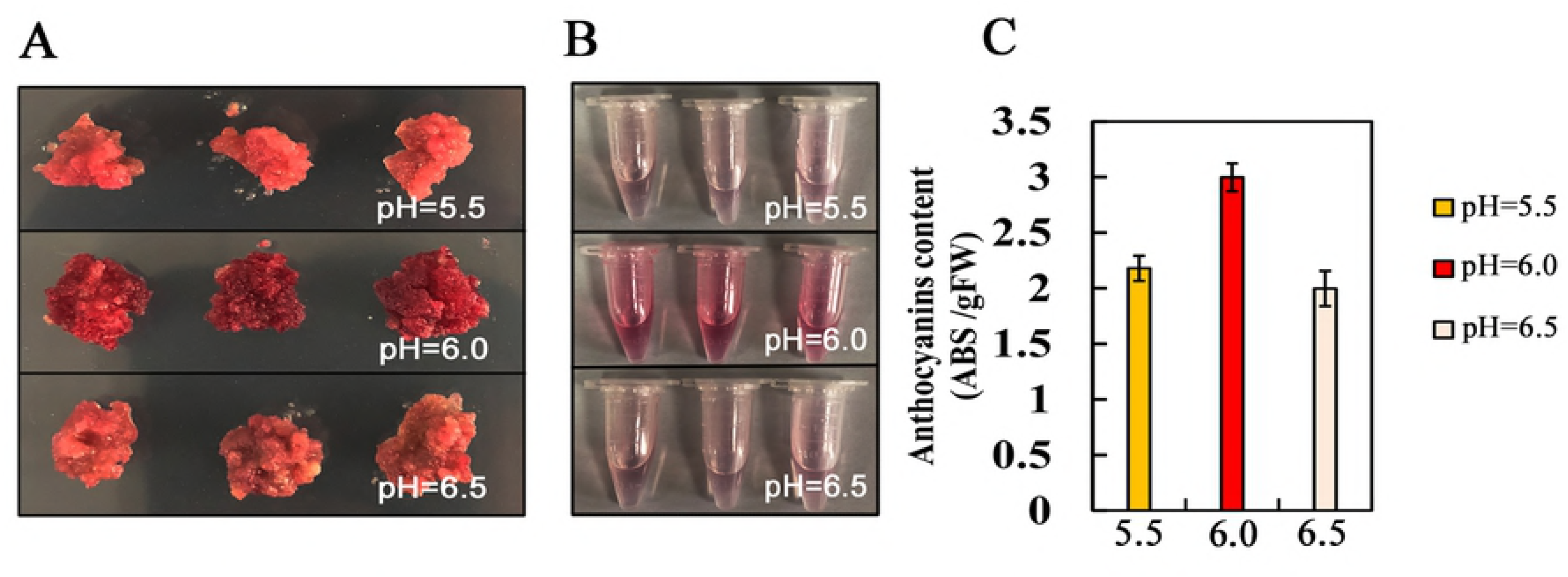
Anthocyanin contents of red-fleshed apple calli exposed to various pH levels (5.5, 6.0, and 6.5). (A and B) Phenotypes of calli treated with different pHs as well as the extraction of anthocyanins. (C) Relative anthocyanin contents over 18 days. The relative anthocyanin content was calculated as follows: absorbance (530 nm)/fresh weight (g).

### Effects of pH on the expression of anthocyanin biosynthesis genes

We investigated whether the changes in anthocyanin contents induced by different pH levels were reflected at the transcript level. The qRT-PCR results revealed that the expression levels of some of the anthocyanin biosynthesis structural genes in red-fleshed apple calli increased with decreasing pH levels, especially *MdCHI, MdDFR*, and *MdUFGT*. Specifically, the *MdDFR* and *MdUFGT* expression levels changed significantly, with the highest transcript levels observed at pH 6.0 (Fig. 2). Furthermore, the *MdMYB3* and *MdMYB10* transcript levels also increased in response to low pH (Fig. 3). Notably, the *MdMYB3* transcript levels increased significantly in calli cultured at pH 6.0 (Fig. 3). The expression levels of these genes were consistent with the changes in anthocyanin content and the degree of pigment precipitation observed at different pHs.

**Fig. 2.**
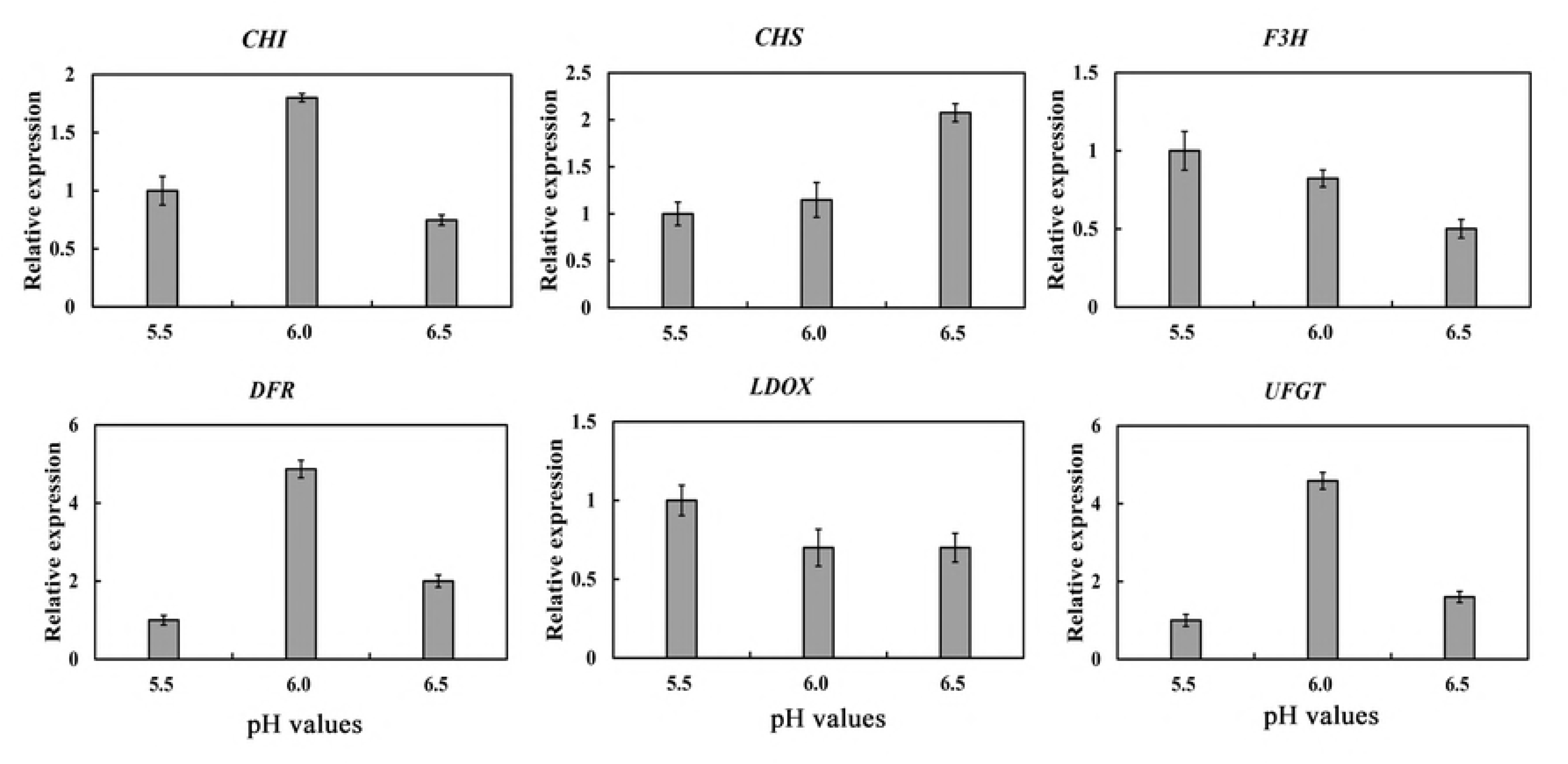
Relative expression levels of anthocyanin biosynthesis structural genes (*MdCHS*, MdCHI, MdF3H, MdDFR, MdLDOX, and *MdUFGT*) in response to different pH treatments. The expression levels for the wild-type (WT) controls grown at pH 5.5 were set to 1.

**Fig. 3.**
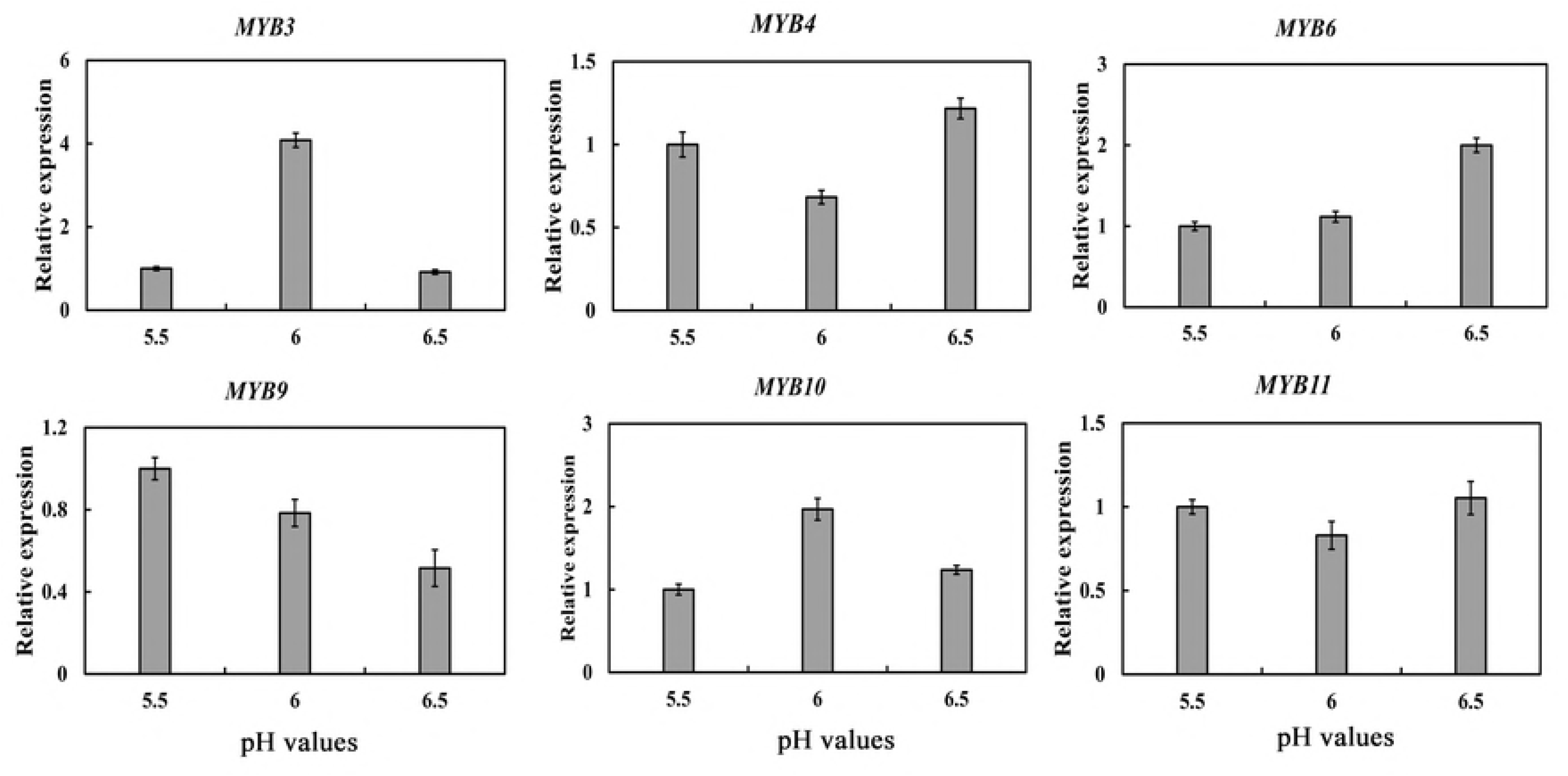
Relative expression levels of anthocyanin regulatory genes (*MdMYB3, MdMYB4, MdMYB6, MdMYB9, MdMYB10*, and *MdMYB11*) in response to different pH treatments. The expression levels for the wild-type (WT) controls grown at pH 5.5 were set to 1.

### *MdMYB3* overexpression enhances anthocyanin biosynthesis in apple calli

The red-fleshed apple calli overexpressing *MdMYB3* changed from red to dark red (Fig. 4A), which may have been associated with the observed significant increase in anthocyanin content (Fig. 4B), when the pH was 5.5, 6.0, and 6.5. The changes in the cyanidin 3-*O*-galactoside content of transgenic calli exposed to various pH levels are described in Table 1. Additionally, the expression levels of structural genes (*MdDFR* and *MdUFGT*) and TF genes (*MdMYB3, MdMYB10*, and *MdMYB11*) were higher in the transgenic calli than in the WT calli (Fig. 5). These results suggested that MdMYB3 promotes anthocyanin accumulation by regulating the expression of genes related to anthocyanin biosynthesis.

**Fig. 4.**
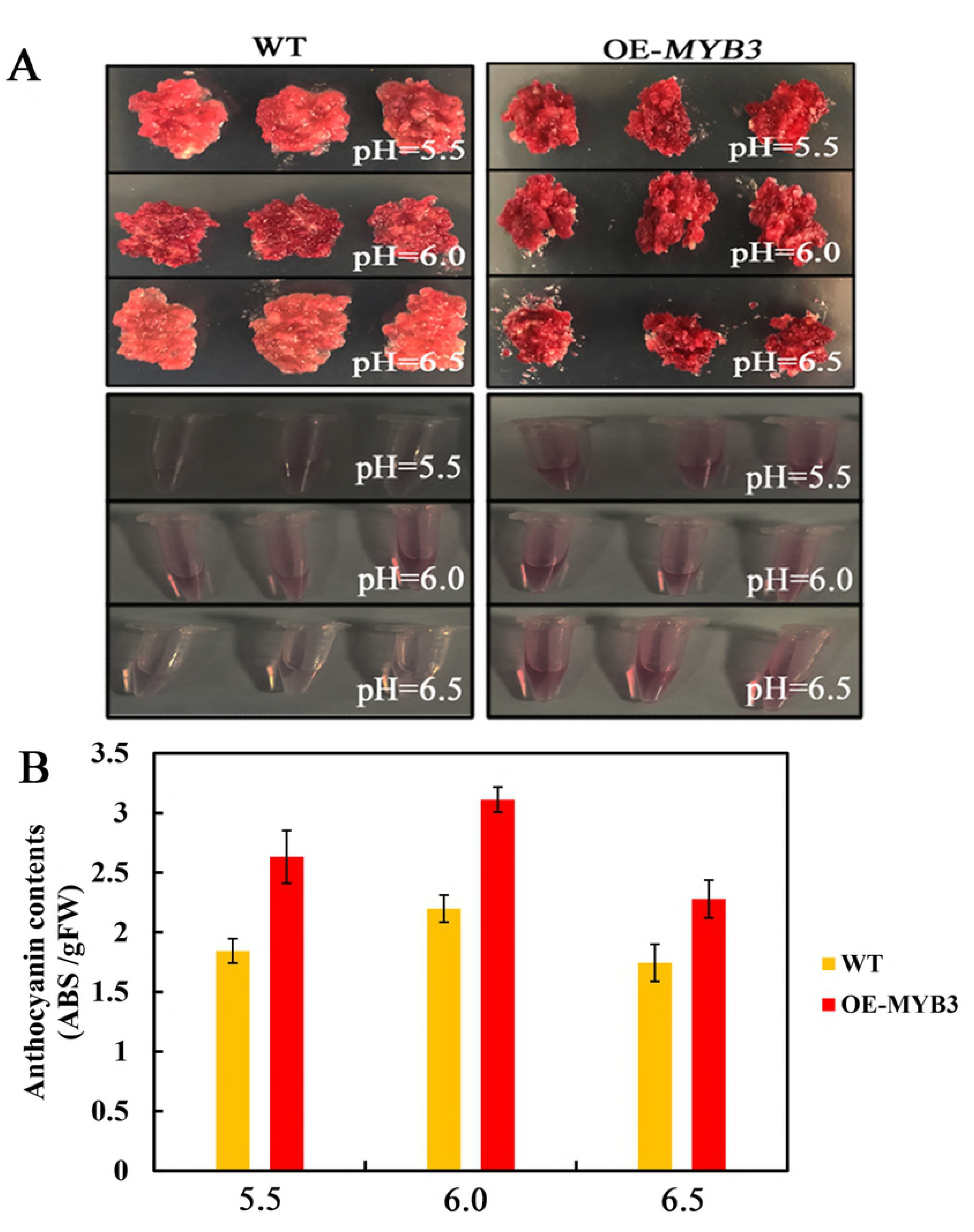
Analysis of transgenic red-fleshed apple calli overexpressing *MdMYB3*. (A) Appearance of the wild-type red-fleshed apple callus and transgenic callus overexpressing *MdMYB3* at different pHs (5.5, 6.0, and 6.5). (B) Anthocyanin levels in the wild-type and transgenic calli at different pHs (5.5, 6.0, and 6.5).

**Fig. 5.**
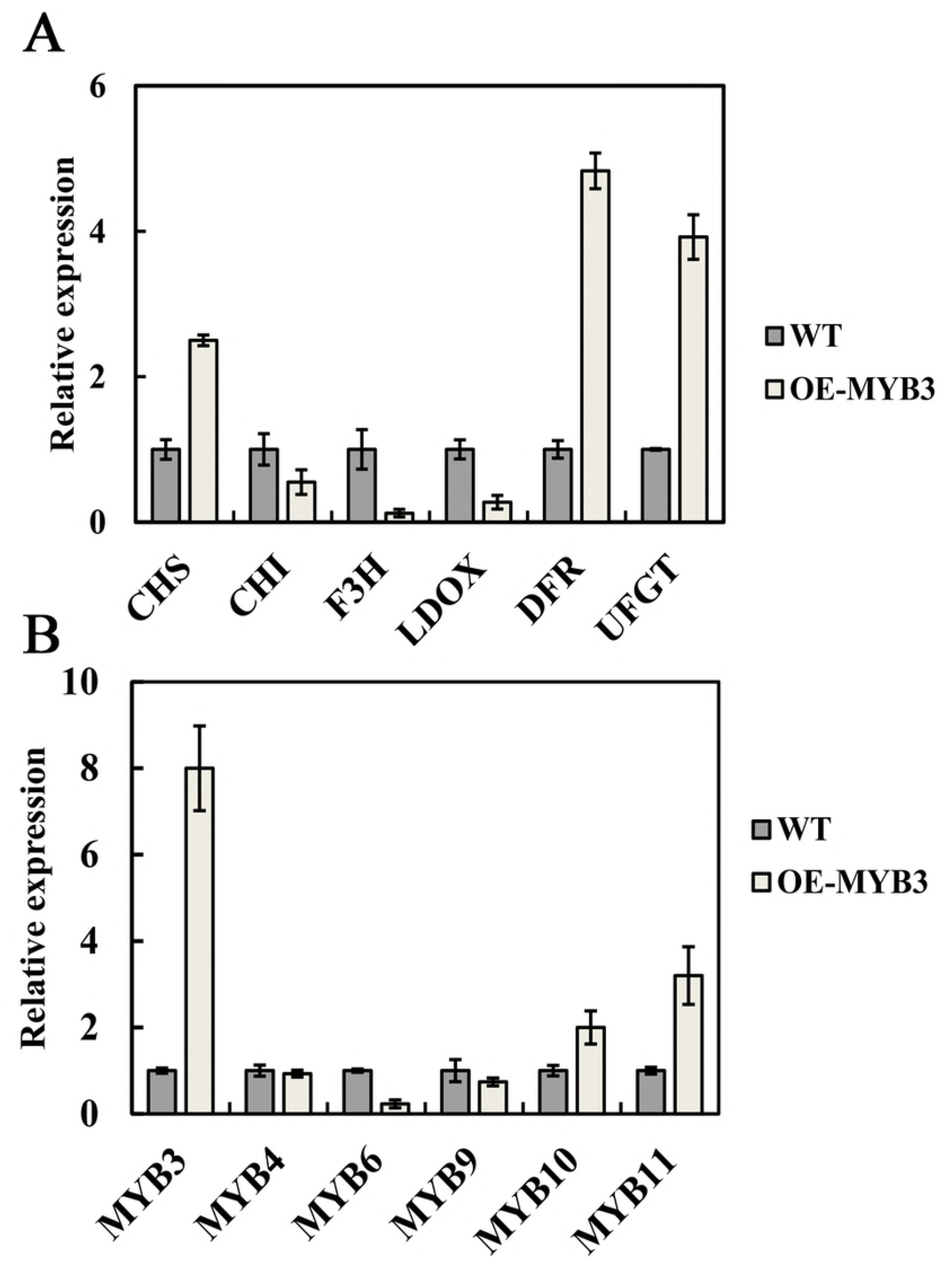
Expression of anthocyanin-related genes in wild-type and transgenic calli at pH 6.0. The expression levels for the wild-type controls were set to 1.

**Table 1.** HPLC data and the cyanidin 3-*O* -galactoside content of each callus. ^a^cyanidin 3-*O* -galactoside standard (Sigma Chemical, St. Louis, MO, USA), WT(PH=5.5): wild-type red-fleshed apple callus at pH 5.5, MdMYB3(PH=5.5): transgenic callus overexpressing *MdMYB3* at pH 5.5, WT(PH=6.0): wild-type red-fleshed apple callus at pH 6.0, MdMYB3(PH=6.0): transgenic callus overexpressing *MdMYB3* at pH 6.0, WT(PH=6.5): wild-type red-fleshed apple callus at pH 6.5, MdMYB3(PH=6.5): transgenic callus overexpressing *MdMYB3* at pH 6.5.

### MdMYB3 directly promotes anthocyanin accumulation

An earlier investigation confirmed that the MYB TFs contain R2R3 domains and play key roles in the anthocyanin biosynthesis pathway [30]. Aligned protein sequences revealed that MdMYB3 contains R2R3 domains. (Fig. 6A). Moreover, the Y1H assay results indicated that MdMYB3 can bind to the *MdDFR* and *MdUFGT* promoters (Fig. 6B). Additionally, we searched the PlantCARE database of *cis-acting* regulatory elements (http://bioinformatics.psb.ugent.be/webtools/plantcare/html/) and determined that the *MdDFR* and *MdUFGT* promoters contain putative MYB-binding site (MBS) elements (Fig. S1). The EMSA results indicated that MdMYB3 can bind to DNA probes containing the *MdDFR* and *MdUFGT* promoter MBS elements, implying that MdMYB3 can specifically recognize these promoters (Fig. 6C). Additionally, our GUS activity analysis indicated that moderately acidic conditions increased the activity of the MdDFR and MdUFGT promoters, and the binding of MdMYB3 to these promoters further enhanced activity (Fig. 6D). Therefore, we speculated that MdMYB3 promotes anthocyanin accumulation by directly regulating the expression of these genes.

**Fig. 6.**
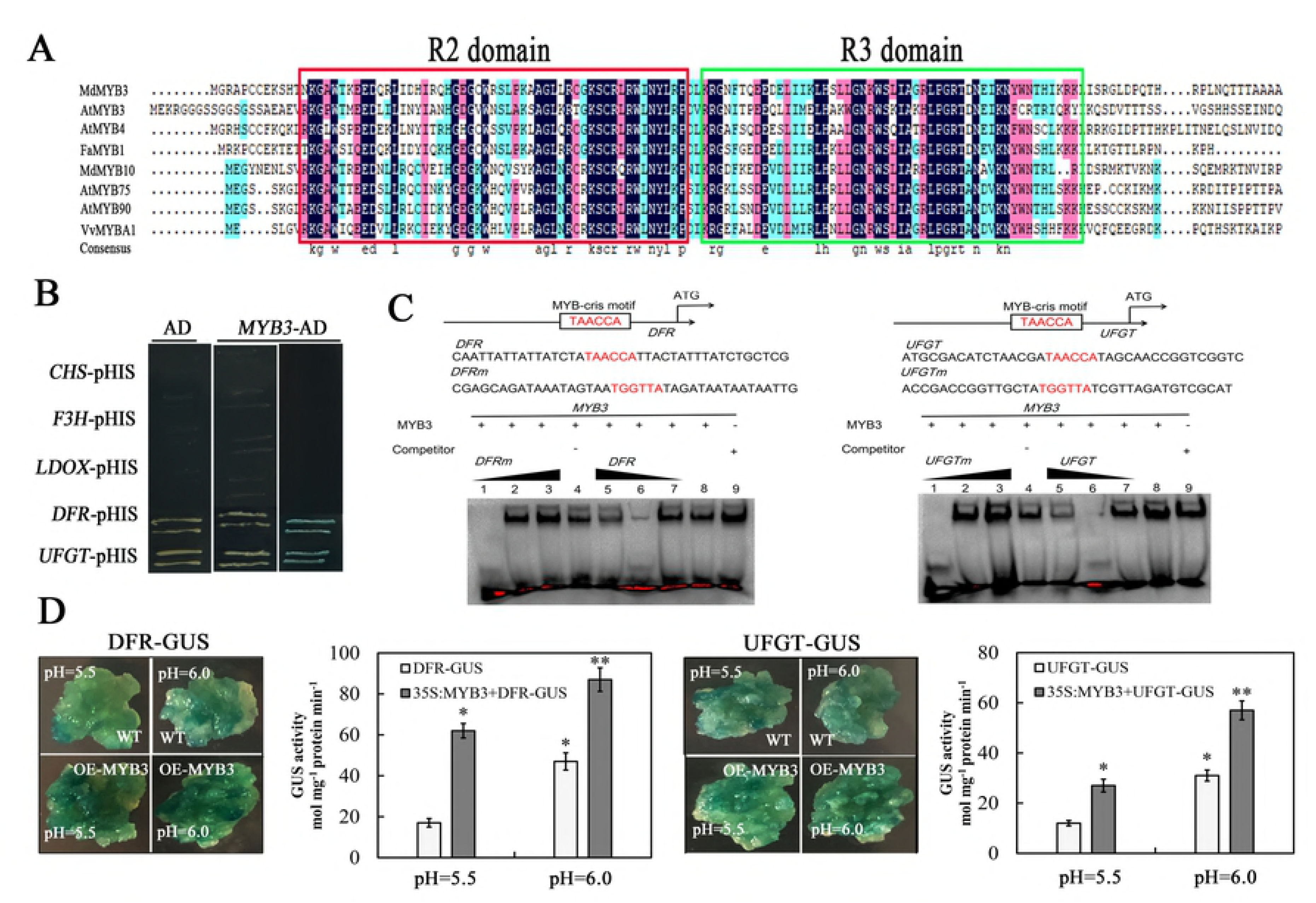
Functional analysis of MdMYB3. (A) Aligned amino acid sequences of the following MYB transcription factors: MdMYB3 (XP_008337875.1), AtMYB3 (AT1G06180), AtMYB4 (AT2G31180), FaMYB1 (AT3G23250), MdMYB10 (XP_009339621.1), VvMYB1 (APM87903.1), AtMYB75 (APM87903.1), and AtMYB90 (APM87903.1). (B) Yeast one-hybrid assay results revealed that MdMYB3 binds to the *MdDFR* and *MdUFGT* promoters. (C) Electrophoretic mobility shift assay results indicated that the MdMYB3-His fusion protein can directly bind to the MYB-binding site of the *MdDFR* and *MdUFGT* promoters *in vitro*. The biotinylated probe was incubated with the MdMYB3-His protein and the freed bound DNA was separated in an acrylamide gel. The unlabeled probe was used as a competitive probe. The 5′-TAACCA-3′ motif was replaced by 5′-GCTACA-3′ in the mutant probe (Mut). (D) The GUS activity assay demonstrated that MdMYB3 activates the *MdDFR* and *MdUFGT* promoters. Transgenic calli producing MdDFR-GUS or MdUFGT-GUS with or without the 35S:MdMYB3 effector were grown and stained at pH 6.0 or 6.5 to detect GUS activity. The mean and standard deviation were calculated using the data from three independent experiments.

## Discussion

Environmental pH is one of the important physical and chemical factors affecting the metabolic activities of plant tissues. Previous studies concluded that a suitable pH can promote anthocyanin biosynthesis in specific tissues of some plant species, including carrot and crabapple [12, 15]. In crabapple seedlings, the synthesis of anthocyanins is beneficial in moderately acidic environments [15]. Consistent with the results of previous studies, we observed that under moderately acidic conditions, especially at pH 6.0, anthocyanins accumulated considerably in apple calli. When the pH of the medium was higher or lower than this threshold, the anthocyanin content decreased. An earlier investigation confirmed that low environmental pH values can promote the accumulation of anthocyanins in crabapple with colored leaves by activating the expression of late anthocyanin biosynthesis genes [15]. In this study, we observed that moderately acidic conditions induced anthocyanin accumulation in red-fleshed apple calli, likely because of changes in the transcription of specific anthocyanin biosynthesis structural genes (*MdDFR* and *MdUFGT*) and a TF gene (*MdMYB3*). Therefore, moderately acidic conditions may promote the accumulation of anthocyanin in apple calli *via* the up-regulated expression of anthocyanin biosynthesis-related genes.

The R2R3 MYB TFs are crucial regulators of anthocyanin biosynthesis and vacuolar pH in plants. In apple, *MdMYB1*, which encodes the R2R3 domain, participates in the regulation of cell pH and anthocyanin biosynthesis [22]. Additionally, another R2R3 MYB TF, MdMYB73, influences malate accumulation and vacuolar acidification in apple [23]. In our study, a protein sequence alignment indicated that MdMYB3 contains the R2R3 domain at the N-terminal. To further investigate the functions of MdMYB3 in red-fleshed apple, we generated transgenic calli overexpressing *MdMYB3* according to an *A. tumefaciens* -mediated transformation method. The anthocyanin contents of the red-fleshed apple calli overexpressing *MdMYB3* increased at the tested pH values. Furthermore, *MdDFR* and *MdUFGT* expression levels were higher in the transgenic calli than in the WT controls, suggesting that MdMYB3 might bind to the promoters of these genes as part of the pH-induced signaling pathways that regulate anthocyanin biosynthesis. Therefore, MdMYB3 may be responsive to changing acidity levels and play a key role in the regulation of anthocyanin biosynthesis.

The *MYB* genes constitute a very large TF family with diverse functions related to many biological processes [29]. The MYB TFs recognize and bind to the MBS in the promoters of the downstream target genes to regulate transcription [30]. Our Y1H assay demonstrated that MdMYB3 binds to the *MdDFR* and *MdUFGT* promoters. Moreover, an analysis involving the PlantCARE database confirmed the presence of multiple MBS sequences in the *MdDFR* and *MdUFGT* promoters. Furthermore, our EMSA data implied that MdDFR and MdUFGT function downstream of MdMYB3. Meanwhile, the GUS activity assay revealed that moderately acidic conditions facilitate the MdMYB3-mediated induction of *MdDFR* and *MdUFGT* expression. Thus, MdMYB3 may be an important regulator of apple coloration in response to acidic conditions.

In conclusion, we revealed that moderately acidic conditions can promote anthocyanin accumulation in apple by up-regulating *MdUFGT* and *MdDFR* expression levels. Furthermore, MdMYB3 helps regulate the coloration of apple plants cultured in media with varying acidity. This regulation is primarily *via* the effects of MdMYB3 on the transcription of the downstream target genes associated with the anthocyanin biosynthesis pathway. The data presented herein may be useful for developing convenient methods for adjusting the coloration of apple fruits that rely on changes to the environmental pH.

## Acknowledgments

This study was supported by grants from the Natural Science Foundation of China (31730080) and the National Key Research and Development Project (2016YFC0501505).

## Author contributions

Conceived and designed the experiments: Yi-Cheng Wang, Jing-Jing Sun, Yan-Ling Wang, and Xue-Sen Chen. Performed the experiments: Yi-Cheng Wang, Jing-Jing Sun, Yan-Fen Qiu, Xiao-Jun Gong, and Li Ma. Analyzed the data: Jing-Jing Sun, Yi-Cheng Wang, Zong-Ying Zhang, and Shou-Qian Feng. Contributed to the writing of the manuscript: Jing-Jing Sun, Yi-Cheng Wang, Yan-Ling Wang, and Xue-Sen Chen. All authors approved the contents of this manuscript.

## Supporting information legends

**Table S1.** Sequences of primers used for PCR validation.

**Fig S1.** Characteristics of the *cis* -acting elements involved in the binding of MYB in the *MdDFR* and *MdUFGT* promoters.

